# Estimating anisotropy directly via neural timeseries

**DOI:** 10.1101/2021.05.25.445605

**Authors:** Erik D. Fagerholm, W.M.C. Foulkes, Yasir Gallero-Salas, Fritjof Helmchen, Rosalyn J. Moran, Karl J. Friston, Robert Leech

## Abstract

An isotropic dynamical system is one that looks the same in every direction, i.e., if we imagine - standing somewhere within an isotropic system, we would not be able to differentiate between different lines of sight. Conversely, anisotropy is a measure of the extent to which a system deviates from perfect isotropy, with larger values indicating greater discrepancies between the structure of the system along its axes. Here, we derive the form of a generalised scalable (mechanically similar) discretized field theoretic Lagrangian that allows for levels of anisotropy to be directly estimated via timeseries of arbitrary dimensionality. We generate synthetic data for both isotropic and anisotropic systems and, by using Bayesian model inversion and reduction, show that we can discriminate between the two datasets – thereby demonstrating proof of principle. We then apply this methodology to murine calcium imaging data collected in rest and task states, showing that anisotropy can be estimated directly from different brain states and cortical regions in an empirical *in vivo* biological setting. We hope that this theoretical foundation, together with the methodology and publicly available MATLAB code, will provide an accessible way for researchers to obtain new insight into the structural organization of neural systems in terms of how scalable neural regions grow – both ontogenetically during the development of an individual organism, as well as phylogenetically across species.

## Introduction

Two of the main concepts upon which computational neuroscience models are based are those of the ‘particle’ [1] and the ‘field’ [2] – both terms that are inherited from theoretical physics.

### Particle theoretic models

In the particle theoretic approach we treat every node within a neural system as a zero-dimensional (point-like) element – a so-called ‘particle’ that evolves in time. The ways in which each one of these neural particles evolve influences the rest of the connected system, such that collectively, the particles form nodes of a dynamically evolving graph [3]. Particle theoretic frameworks yield experimental advantages for neuroimaging modalities such as electroencephalography (EEG), in which there are usually very few measurement locations. Furthermore, particle theoretic frameworks have computational and statistical advantages for neuroimaging analyses due to associated dimensionality reduction – an attribute that becomes increasingly important for large-scale recordings of neural systems [4]. However, this computational expediency comes at the cost of losing the spatial information contained in a continuum description.

### Field theoretic models

On the other hand, the field theoretic approach treats a neural system as a continuous structure called a ‘field’ that is a function of position, with position treated as a continuous variable. A neural field can exist in 2D or 3D space: it is natural to work in a twodimensional space when modelling a single cortical sheet or a three-dimensional space for a cross-cortical volume [5]. In this paper, we use a model that is in essence a compromise between the particle and field models, by taking a continuous field and discretizing it such that we only consider certain points in space – which we henceforth refer to as a discretized field theoretic model.

### Isotropic vs. anisotropic systems

A system is said to be isotropic if it looks identical in every direction. This means that if we imagine ourselves standing somewhere within an isotropic structure, then we would see precisely the same structure along all lines of sight. Conversely, anisotropy is a measure of the extent to which a system deviates from perfect isotropy. For example, a sheet of wood is anisotropic due to the preferential directionality of the grain – which we can see by the fact that it is easier to break the wood along the grain than it is to break it against the grain. We present a discretized field theoretic model that allows for the estimation of anisotropy in connected dynamical systems of arbitrary dimensionality. We provide accompanying MATLAB code in a public repository that can be readily used to measure levels of anisotropy on a node-wise basis via timeseries measurements.

### Overview

This paper comprises three sections.

In the first, we outline the theoretical foundations of Lagrangian field theory and the form of a generalised scalable discretized equation of motion that can be used both for forward generative models and for model inversion via neural timeseries.

In the second section, we generate *in silico* data via forward models of an isotropic and an anisotropic system. We then use Bayesian model inversion and subsequent Bayesian model reduction to show that we can correctly discriminate between the isotropic and anisotropic systems – thereby providing construct validation.

In the third section, we use murine wide-field calcium imaging data [6–8] collected in both rest and task states to map levels of anisotropy across different cortical regions directly via the *in vivo* timeseries.

We suggest that the presented methodology could be valuable in future large-scale studies of neural systems, in which the quantification of region-wise anisotropy may shed light on how neural systems grow both phylogenetically within the lifespan of an individual animal, as well as ontogenetically across species [9].

## Methods

We will now cover the technical foundations of the approach, starting with Lagrangian field theory and the principle of stationary action. We then derive a generalised, scalable, discretized field theoretic Lagrangian and consider the empirical estimation of anisotropy under this formulation using empirical (timeseries) data and Bayesian estimators.

### Lagrangian field theory

We remind the reader of the basic concepts underlying Lagrangian field theory and the principle of stationary action in Appendix I. In brief: we represent the state of a system by a field which is a function of the 4D space-time position *r* ≡ (*t, x, y, z*). The equations of motion that describe how this field evolves in time are obtained by requiring that the field *φ*(*r*) renders the value of a quantity known as the action S stationary:

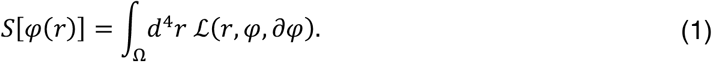

The integral in the definition of the action is over the space-time domain Ω encompassing all space from the initial time *t_i_* to the final time *t_f_*. The integrand, which is known as the Lagrangian density 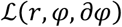, defines the system of interest as a function of *r*, the values of the fields *φ* at *r*, and their spatiotemporal derivatives *∂_φ_* at *r*.

### Scale transformations

We define a scale transformation as a mapping from arbitrary points in the 5-D space with axes labelled (*φ, r*) = (*φ, t, x, y, z*) to scaled points (*φ_s_, r_s_*) = (*λ_φ_, λ^a^r*), where *λ* is an arbitrary scale factor and *α* is a constant. A field configuration *φ* = *φ*(*r*) is a 4-D surface in this 5-D space, and the scale transformation takes points on that surface to points on a new 4D surface, defining a new field configuration. The value of the new field *φ_s_* at the scaled spacetime point *r_s_* = *λ^a^r* is related to the value of the original field *φ* at the unscaled point *r* via

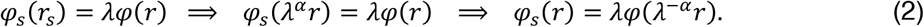

It is convenient to allow different scaling exponents in different space-time directions, so from now on *λ^a^r* is to be understood as shorthand for the vector (*λ^a_t_^t, λ^a_x_^x, λ^a_y_^y, λ^a_z_^z*).

Taking partial derivatives of Eq. (2), we obtain:

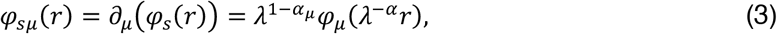

where λ^1−*α*_*μ*_^ depends only on the *μ*^th^ component of the vector of exponents *α* = (*α_t_, α_x_, α_y_, α_z_*).

From now on, we denote the vector with components λ^1−*α*_*μ*_^ *φ_μ_*(*λ^−α^r*) as *λ*^1−*α*^*∂*_*φ*_(*λ*^−*α*^*r*).

### Scaling the action

Using Eqs. (39), (2) and (3), we see that the scaled action is given by:

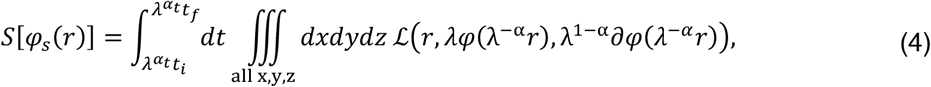

We then change variables in Eq. (4), setting *r*′ = *λ^-α^r* such that:

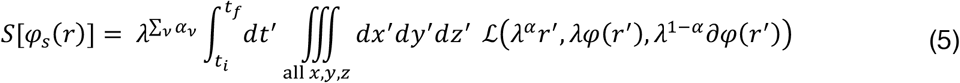

where *λ*^∑_*v*_*α*_*v*_^ is the Jacobian that accounts for the change in integration variables and ∑_*v*_ *α_v_* = *α*_t_ + *α_x_* + *α_y_* + *α_z_*. The integrals are now over the same space-time region Ω as in the original un-scaled action in Eq. (39), which means that we can re-write Eq. (5) using the same simple notation:

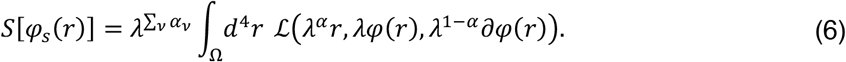

### Scalable systems

The action *S*[*φ*(*r*)] is said to be scalable, or equivalently to possess ‘mechanical similarity’ [10] if, for any choice of *φ*(*r*), not just choices that make the action stationary and therefore satisfy the Euler-Lagrange equation of motion (see Appendix I), the following relationship holds:

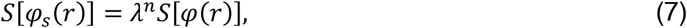

where *n* is a constant. Note that a ‘scalable’ system should not be confused with a ‘scale free’ system, which is one that lacks a characteristic length scale, such as those studied in the physics of phase transitions.

More explicitly, we can use Eqs. (6) and (7) to express the scalability condition as follows:

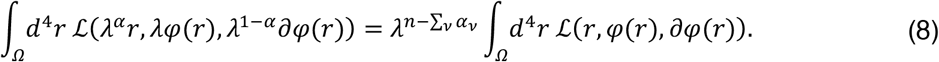

### Generalised scalable Lagrangians

We can expand any analytic Lagrangian density 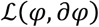 as a power series:

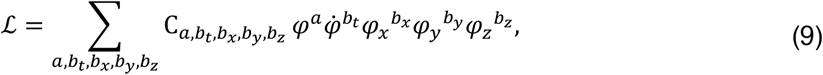

where C*_a, b_t_, b_x_, b_y_, b_z__* is an expansion coefficient and the summations over the integers *a, b_t_, b_x_, b_y_, b_z_* range from 0 t_o_ ∞. We have assumed for the sake of simplicity that the Lagrangian density has no explicit dependence on *r*. This is normally the case when the system of interest is not driven by external forces or other influences. We next use Eq. (9) to obtain 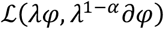:

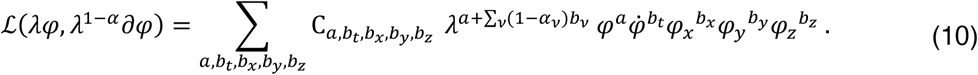

For the action to be scalable, Eq. (8) tells us that Eq. (10) must equal:

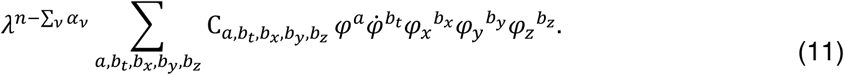

We conclude that the action is scalable if and only if

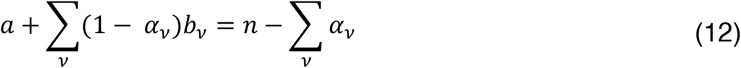

for all choices of the integers *a* and *b_v_* at which C*_a, b_t_, b_x_, b_y_, b_z__* is non-zero. If, for example, we consider possible contributions to the Lagrangian with specific values of *b_t_*, *b_x_*, *b_y_*, and *b_z_*, Eq. (12) tells us that C*_a, b_t_, b_x_, b_y_, b_z__* must be zero unless *α* = *n* – ∑_*v*_ *α_v_* – ∑_*v*_(1 – *α_v_*)*b_v_*. The value of *α* is determined by the values of *b_t_, b_x_*, *b_y_*, *b_z_* and the summation over *a* is no longer required. The generalised scalable discretized field theoretic Lagrangian may therefore be written as follows:

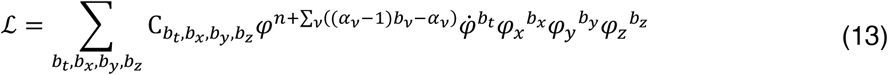

### The anisotropic wave equation

Let us now design a special case of Eq. (13) in two spatial dimensions. Having chosen to set *α_t_* = *α_x_* and *n* = *α_y_* + 2, we construct a Lagrangian density with three non-zero terms. In the first term, C*_b_t_, b_x_, b_y__* = 1, *b_t_* = 2, and *b_x_* = *b_y_* = *b_z_* = 0; in the second term, C*_b_t_, b_x_, b_y__* = −1, b_x_ = 2, and *b_t_* = *b_y_* = *b_z_* = 0; and finally, in the third term, C*_b_t_, b_x_, b_y__* = −1, *b_y_* = 2, and *b_t_* = *b_x_* = *b_z_* = 0.

This yields the Lagrangian density:

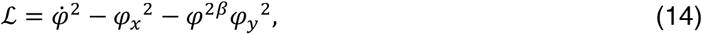

where the exponent *β*, which is by defined by *β* = *α_y_* – *α_x_*, quantifies the degree of anisotropy, such that the system is perfectly isotropic when *β* = 0 ⇒ *α_y_* = *α_x_*. The corresponding equation of motion – the two-dimensional Euler-Lagrange equation (see Appendix I) – is:

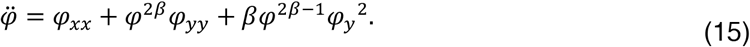

We can verify that if *φ*(*t, x, y*) is a solution of this equation, so is the scaled field 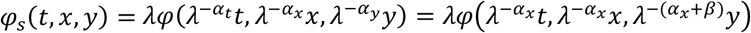 (see Appendix II).

In the case of an isotropic system, when *β* = 0, the Euler-Lagrange equation becomes:

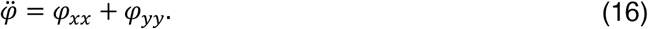

We see that the Lagrangian density of Eq. (14) leads to an equation of motion (15) that reduces to the wave equation (16) in the case of an isotropic system. This makes it an intuitive test case.

### Discretisation and noise

For Eq. (15) to be used in the modelling of neural timeseries we must first discretize the partial spatial derivatives. This is necessary because we do not deal with spatially continuous data in neuroimaging, but rather with data collected at a discrete set of points. We therefore make the following standard transformations:

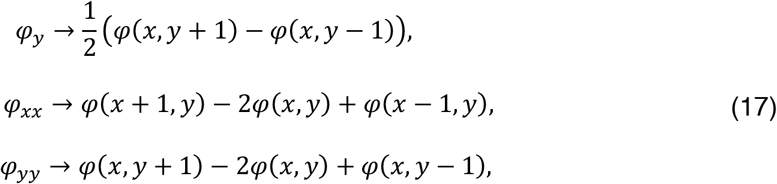

where *φ*(*x, y*) is the value of the field at the point (*x, y*) and, e.g., *φ*(*x* + 1, *y*) is the value of the field one ‘step’ in the positive *x* direction in the graph from the point (*x, y*).

Applying the transformations to Eq. (15), we obtain:

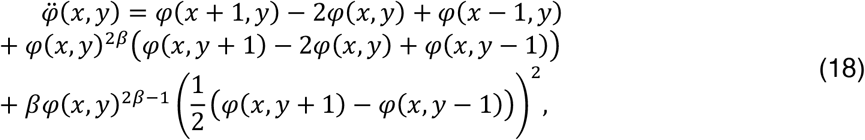

which we split into two first-order differential equations by defining a new variable to obtain the final form of the equations of motion used in all subsequent forward models and Bayesian model inversions presented in this paper:

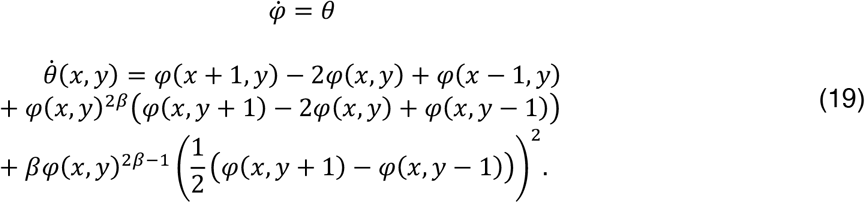

where the observation equation consists of the *φ* variable.

We then use Eq. (19) as the equations of motion for subsequent model inversion with the Statistical Parametric Mapping (SPM) software. The associated observer equation comprises the *φ* variables – i.e., we assume that the strength of the field *φ*(*x, y*) is what is being measured in the neural timeseries at the (*x, y*) coordinate.

Eq. (19) is the basis of the MATLAB code made available for the use with forward generative models, as well as with Bayesian model inversion of timeseries of arbitrary dimensionality from any neuroimaging modality.

### Synthetic data

We consider a 2D grid of size 3×3 pixels, where each of the nine pixels is given different initial conditions and subsequently allowed to evolve according to the equation of motion in Eq. (19). We run two forward models: a) one isotropic case in which *β* = 0; and b) one anisotropic case in which *β* ≠ 0. Having created these synthetic data with a prior of *β* = 0, we then perform Bayesian model inversion using Dynamic Expectation Maximization (DEM) [11] to infer the latent states and estimate both the *β* parameter, as well as the hyperparameters – the components of precision of random fluctuations on the observation noise and states. Model inversion for any discrete system can be performed on a node-by-node basis, by considering the ways in which the dynamics evolve in the immediate neighbourhood of the current node under consideration. When this model is equipped with fluctuations one can use standard (Variational Laplace) Bayesian model inversion procedures to estimate the exponents for any given timeseries. We then set the prior for the free parameter *β* to zero and DEM to obtain a posterior estimate for *β* from both the synthetic isotropic and anisotropic data. Following model inversion, we use Bayesian model reduction [12, 13] to test the evidence for a perfectly isotropic system in which *β* = 0 by setting the prior variances of *β* to zero. We are therefore able to test whether we can correctly identify the ground truth isotropic data (created with *β* = 0) with a higher evidence for the reduced model and conversely whether we can correctly identify the ground truth anisotropic data (created with *β* ≠ 0) with a higher evidence for the full model.

### Empirical data

All animal experiments were carried out according to the guidelines of the Veterinary Office of Switzerland following approval by the Cantonal Veterinary Office in Zürich. All murine calcium imaging data were collected as previously reported [6–8]. As with the synthetic data, we perform Bayesian model inversion to obtain posterior estimates for the *β* parameter quantifying the extent to which the time series for each pixel deviate from isotropy at *β* = 0. We perform this model inversion once for every second pixel (*n* = 6651) within each trial (*n* = 10), mouse (*n* = 3) and condition (*n* = 2, task and rest). Following model inversion, we average the posteriors for the *β* parameter across trials to obtain results per mouse and condition and then average these posteriors once more across mice. We then filter these averaged images with 2-D Gaussian moving average smoothing kernels with a semi-width window of 7 pixels.

## Results

We show the ways in which the synthetic timeseries evolve for the isotropic (Figure 1A) and anisotropic (Figure 1B) cases. Following model inversion and reduction, we then demonstrate proof of principle by showing that there is higher evidence for the ground truth isotropic data having been created with the isotropic model (Figure 1C) and conversely for the ground truth anisotropic data having been created with the anisotropic model (Figure 1D). Finally, we show the different degrees of anisotropy in the murine calcium imaging data in rest and task states (Figure 1E). Overall, there is a marked variability in the degrees of anisotropy across mice and states. On the other hand, the secondary motor cortices show consistently high degrees of anisotropy across mice and states. The generation (Figure 1 A&B) and inversion (Figure C & D) for the synthetic data can be fully reproduced with the accompanying MATLAB code and the murine calcium imaging data in Figure 1 E & F is made available in a public repository.

**Figure 1:**
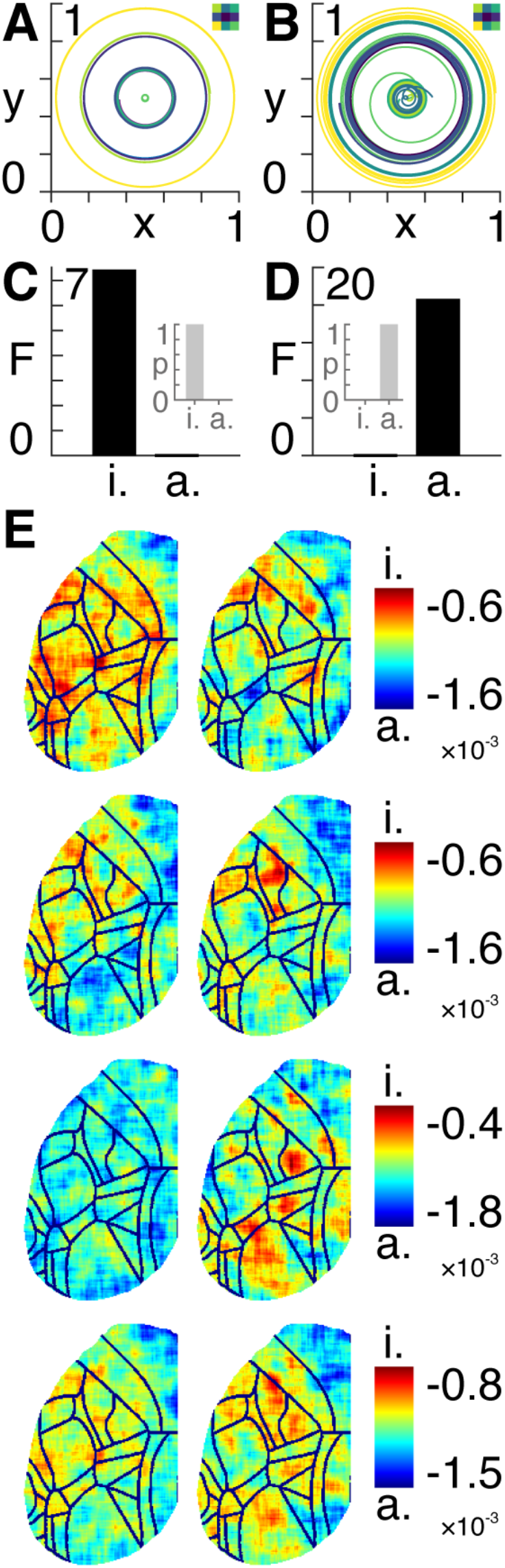
Synthetic and experimental data. **A)** Synthetic data generated using the isotropic model in Eq. *(19)* with β = *0*. The colours of the wavefronts correspond to corresponding pixel in the grid inset top right. The x and y axes show the amplitudes of the wavefronts multiplied by cos(time) and sin(time), respectively. **B)** Synthetic data generated using the anisotropic model in Eq. *(19)* with β ≠ *0*. The colours of the wavefronts correspond to corresponding pixel in the grid inset top right. The x and y axes show the amplitudes of the wavefronts multiplied by cos(time) and sin(time), respectively. **C)** Approximate lower bound log model evidence given by the free energy (F) following Bayesian model reduction for isotropic (i.) and anisotropic (a.) models using the isotropic ground-truth data. Corresponding probabilities (p) derived from the log evidence are shown in the inset on the right. **D)** Approximate lower bound log model evidence given by the free energy (F) following Bayesian model reduction for isotropic (i.) and anisotropic (a.) models using the anisotropic ground-truth data. Corresponding probabilities (p) derived from the log evidence are shown in the inset on the left. **E)** Left hemisphere of calcium imaging data collected in three mice (first three rows) in rest (left column) and task (right column) states. The final fourth row shows average values across the three mice. The colour bars indicate the value of the β exponent ranging from isotropic (i.) to increasingly anisotropic (a.) pixels.

## Discussion

We present a theoretical framework, together with a practical numerical analysis, designed for the estimation of anisotropy in arbitrary timeseries from any connected dynamical system. The basis for this framework rests upon classical Lagrangian field theory applied to scalable (mechanically similar) dynamical systems. Scalability entails a situation whereby a system continues to obey the same equation of motion as it changes size. In other words, a dynamical system that grows or shrinks will begin producing states that are different from those of the original (unscaled) system. However, in systems possessing the property of scalability, the new states in the scaled system will still be described by the same equation of motion used to describe the original (unscaled) system.

It stands to reason that the dynamical evolution of the signals propagating in neural systems possess some form of scalability, given that evolutionary processes add new neuroanatomy to existing structures i.e., the same basic architecture extends itself whist maintaining information processing capabilities [14–16]. Similarly, a neural structure changes scale during development, whilst allowing for information processing to remain intact. It is therefore of interest to develop tools that allow for the analysis of scalable systems. We therefore pose the following question: given that neural systems possess some form of scalability, what are the consequences thereof and what further questions present themselves? It is in this spirit that we present a formalism that applies to any scalable dynamical system that is sufficiently general to accommodate the evolution of any (driven or non-driven) system in any number of spatial dimensions.

With reference to the murine calcium imaging results, we note the following three main results: a) there is a high variability in anisotropy across mice and states, which may be due to the low number of trials and/or to the fact that anisotropy values may vary from trial to trial depending on neural activity; b) there is no clear difference in anisotropy across rest/task states; and c) there are consistently high levels of anisotropy in the secondary motor cortex (Mos) across mice and states, which may reflect the non-homogeneous nature of local networks.

We construct a methodology that is set within the Bayesian model inversion scheme of Dynamic Causal Modelling (DCM). This means that, by using the timeseries measured from any (e.g., neuroimaging) modality, we obtain posterior estimates of spatial stretch factors – one for each spatial dimension. The discrepancy between these stretch factors then directly provides an estimate of the anisotropy at every voxel in the neuroimaging data. We thus obtain an intuitive understanding of what these measures mean by imagining ourselves standing at a certain node in a neural system. If the system is isotropic then, as we look in every direction – up-down, leftright, and back-forward, we will see no difference in the ways in which the signals evolve in time in these different directions. On the other hand, if the system is anisotropic then we will see a difference in our lines of sight along the different axes – and the greater this difference becomes the greater the degree of anisotropy. We demonstrate proof of principle by generating synthetic data using a special case of the generalised Lagrangian that reduces to the wave equation in the limit of the perfectly isotropic case. Using Bayesian model inversion and reduction, we show that we can correctly identify which of the two models (anisotropic and isotropic) were used to generate each dataset, thus showing the discriminatory ability of the proposed methodology.

An empirically determined estimation of anisotropy could be informative in imaging neuroscience, as it facilitates a direct empirical measurement of how sub-structures within the brain grow (under the assumption of scalability). This provides a new way of assessing the ways in which anatomy changes across both an evolutionary timeline, as well as across the lifespan of an individual organism. It is our hope that the theory, methodology, and accompanying tools will allow for these kinds of questions to be addressed by researchers and that these will lead to a clearer understanding of spatial dependencies, growth, and development of neural systems.

## Appendix I

The classical real scalar field of interest in this work depends on position and time, and it turns out to be convenient to treat it as a function of the four-dimensional Cartesian vector

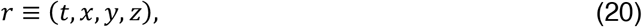

where *t* is time, and *x, y, z* are spatial length, width, and height coordinates, respectively. Individual components of the vector *r* are written *r_μ_*, with *μ* any element of the set {*t, x, y, z*}. The field *φ* is a function of *r*:

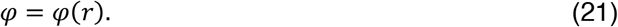

The vector *∂_φ_* of partial derivatives of *φ* at *r* is given by

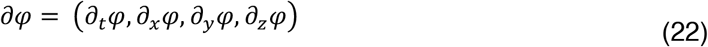

and its components are written *∂_μ_φ* or, more simply, *φ_μ_*.

The central quantity in Lagrangian field theory is the Lagrangian density, 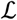, which is a function of *r, φ*, and *∂_φ_*:

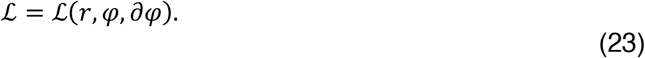

Note that we have not yet assumed any relationship between the values of *r, φ* and *∂_φ_*; the Lagrangian density can be evaluated for any choices of the 9 real numbers required to specify the scalar field *φ* and the two four-component vectors *r_μ_* and *∂_μ_φ*.

Given a particular choice of field ‘trajectory’ *φ*(*r*), the standard definition of the action as a functional of *φ*(*r*) is:

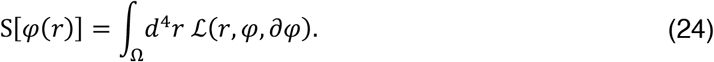

A trajectory in this context consists of the values of the field *φ*(*r*) = *φ*(*t, x, y, z*) at all spatial points (*x, y, z*) and all times *t* between a chosen initial time *t_f_* and a chosen final time *t_f_*. The four-dimensional integration volume *Ω* coincides with the region in which the trajectory is defined. Note that we are now assuming that the field *φ* and its derivatives *φ_μ_* are functions of *r*, so *φ* and *∂_φ_* are now related to each other. We are also assuming that *φ* and *φ_μ_* tend to zero as the spatial distance 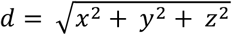 from the origin tends to infinity.

The principle of stationary action tells us that the evolution of *φ*(*r*) between the initial and final times, *t_i_* and *t_f_*, renders the action stationary with respect to all variations

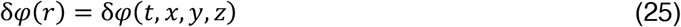

that vanish when *t* = *t_i_* and *t* = *t_f_*.

Using Eq. (24), we evaluate the variation of the action as follows:

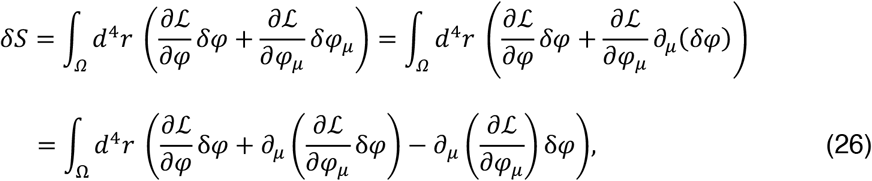

where we are using the Einstein summation convention according to which terms in which the same suffix appears twice are automatically summed over all four values of that suffix. We next convert the middle term on the second line to a surface integral by using the 4-D version of the divergence theorem to obtain:

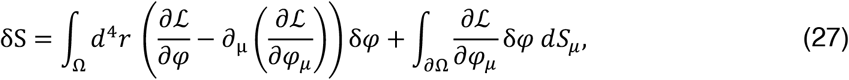

where *∂Ω* is the surface of the 4-D volume *Ω* and *dS_μ_* is an element of the 3-D surface of Ω. If the field decays to zero rapidly enough as the spatial distance from the origin tends to infinity, and remembering that *δ_φ_* = 0 when *t* = *t_i_* and *t* = *t_f_*, the surface integral vanishes, and we obtain:

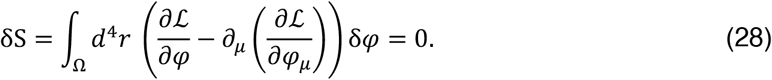

Since *δ_φ_* is arbitrary except for the constraint that it vanishes at the surface *∂Ω*, the principle of stationary action (*δS* = 0) implies that the fields evolve according to the Euler-Lagrange equation:

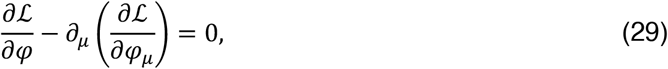

or more explicitly:

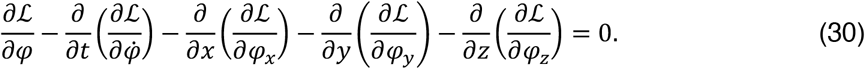

## Appendix II

To show that the scaled field *φ_s_* = *λ_φ_*(*λ*^-*α*_*x*_^*t, λ*^−*α*_*x*_^*x, λ*^−(*α_x_*+*β*)^*y*) is also a solution of Eq. (15), we differentiate *φ_s_* twice with respect to time:

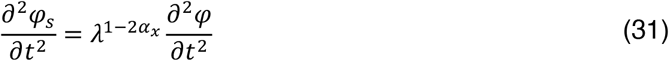

as well as twice with respect to the *x* coordinate:

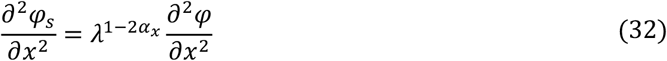

and twice with respect to the *y* coordinate:

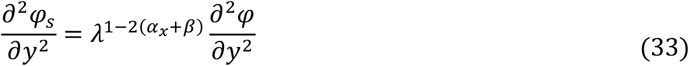

We then take raise *φ_s_* to the power of 2*β*:

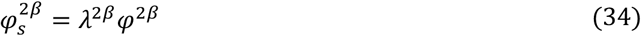

which, together with Eq. (33), means that:

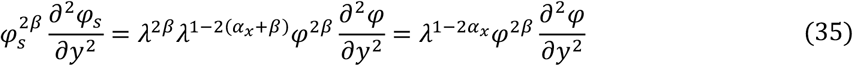

We then differentiate *φ_s_* once with respect to the *y* coordinate and take the square:

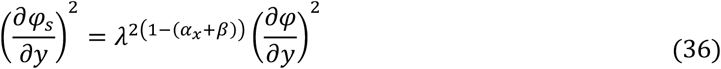

which, together with Eq. (34), means that:

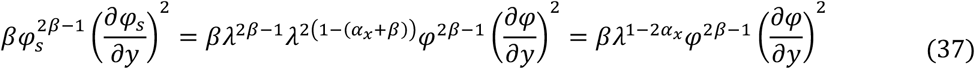

Therefore, by using Eqs. (31), (35) and (39) with the original equation of motion in Eq. (15):

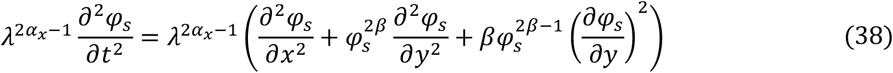

where the *λ*^2*α*_*x*_−1^ factor cancels, leaving the form of the original equation of motion in Eq. (15):

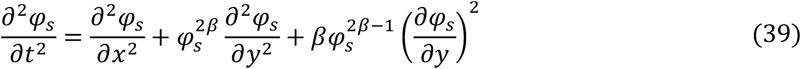

We have therefore shown that if *φ*(*t, x, y*) is a solution of Eq. (15), so is the scaled field 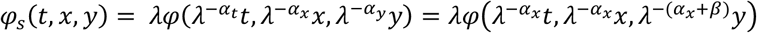.

## Data availability

The MATLAB code used to produce Figure 1 A, B, C & D is made available at the following public repository: github.com/allavailablepubliccode/anisotropy

All pre-processed murine calcium imaging data used to produce Figure 1 E & F are made available in the following public repository: figshare.com/articles/Murine_calcium_imaging_data/12012852

## Author contributions

All authors conceived of the analysis and wrote the paper.

## Acknowledgements

E.D.F. was supported by a King’s College London Prize Fellowship; K.J.F. was funded by a Wellcome Principal Research Fellowship (Ref: 088130/Z/09/Z); R.J.M was funded by the Wellcome/EPSRC Centre for Medical Engineering (Ref: WT 203148/Z/16/Z); R.L was funded by the MRC (Ref: MR/R005370/1). The authors would also like to acknowledge support from the Data to Early Diagnosis and Precision Medicine Industrial Strategy Challenge Fund, UK Research and Innovation (UKRI), the National Institute for Health Research (NIHR), the Biomedical Research Centre at South London, the Maudsley NHS Foundation Trust, and King’s College London.

